# Impairment of the blood brain barrier accelerates a negative ultraslow potential in the locust CNS

**DOI:** 10.1101/2025.04.11.648410

**Authors:** R. Meldrum Robertson, Andrew Donini, Yuyang Wang

## Abstract

Insects provide useful models for investigating evolutionarily conserved mechanisms underlying electrical events associated with brain injury and death. Spreading depolarizations (SD) are transient events that propagate through neuropil whereas the negative ultraslow potential (NUP) is sustained and reflects accumulating damage in the tissue. We used the locust, *Locusta migratoria*, to investigate ion homeostasis at the blood brain barrier (BBB) during SD and NUP induced by treatment with the Na^+^/K^+^-ATPase inhibitor, ouabain. We found that sustained SD caused by the metabolic inhibitor, sodium azide, was associated with a large reduction of K^+^ efflux through the BBB at ganglia (= grey matter) but not at connectives (= white matter). This was accompanied by a large increase in tissue resistivity but no conductance changes of identified motoneuron dendrites in the neuropil. Males recovered more slowly from ouabain-induced SD, as previously described for anoxic SD. Impairment of barrier functions of the BBB pharmacologically with cyclosporin A or DIDS, or by cutting nerve roots, accelerated the NUP, thus promoting earlier and more frequent SD, but had no effect on the temporal parameters of SD. We conclude that the mechanisms underlying onset and recovery of SD are minimally affected by the damage associated with the NUP. We suggest that future research using tissue-specific genetic approaches in *Drosophila* to target identified molecular structures of the BBB are likely to be fruitful.

**New and Noteworthy:** Inhibition of the sodium pump in the locust CNS causes repetitive spreading depolarization (SD) and a negative ultraslow potential (NUP) providing a model for investigation of phenomena relevant to human health. We show that impairment of the blood brain barrier accelerates the NUP but has no impact on the trajectory of SD events. Hence, rapid mechanisms of onset and recovery of ion homeostasis occur against a background of slowly increasing neural damage.

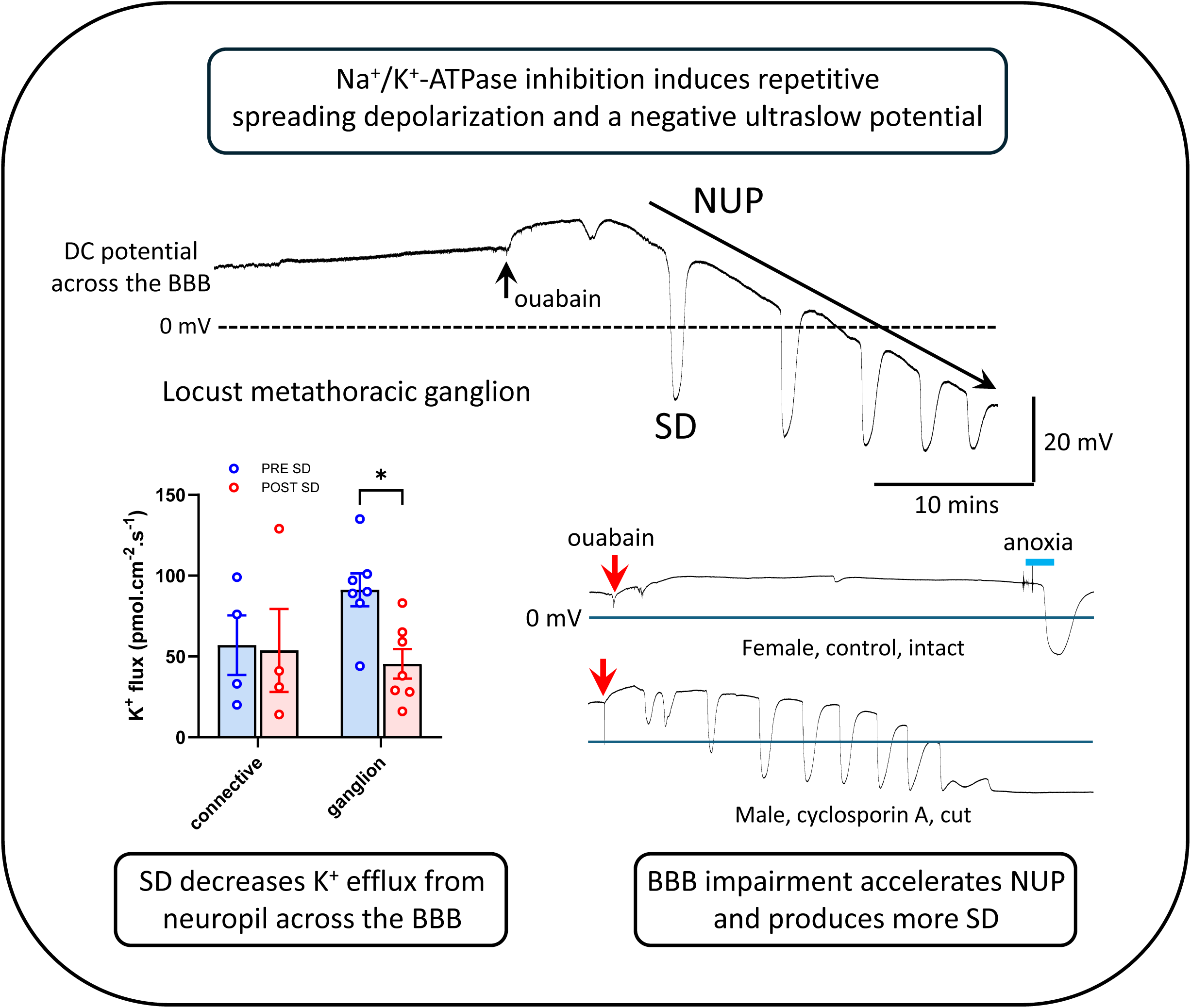

## Introduction

Brain death in mammals, including humans, is characterized by a progression of electrical events in the tissue: an initial non-spreading depression of signalling activity is followed by transient, reversible spreading depolarisation (SD) events, and a sustained, terminal SD that often leads into a large, negative ultraslow potential (NUP) (Charpier, 2023; Dreier et al., 2025; Drenckhahn et al., 2012; Lückl et al., 2018; Sword et al., 2025). At a critical point in this sequence, depending on energetic resources in the tissue, recovery is impossible, cells die, and damage accumulates thereby generating the NUP. Since its initial description (Leão, 1947; Leão, 1944), much has been learned about SD and its consequences (Andrew et al., 2022; Dreier and Reiffurth, 2015; Lemale et al., 2022), however the cellular and molecular mechanisms underlying these events are not completely understood.

Across diverse species in the animal kingdom, SD has been clearly identified in vertebrates and insects, where it is associated with rapid information processing underlying complex behaviours enabled by tight regulation of the extracellular ionic environment of neuropil by virtue of energetically expensive blood brain barriers (BBB) (Robertson et al., 2020). A detailed phylogenetic analysis shows that the convergent evolutions of BBBs and susceptibility to SD completely overlap suggesting that SD is an emergent property of nervous systems adapted for speedy ionic and osmotic control of a restricted interstitium (Andersen et al., 2025). The widespread occurrence of SD lends itself to a comparative experimental approach and the benefits of using model organisms to generate testable hypotheses are well known (e.g. (Spong et al., 2017)).

Insects, particularly fruit flies and locusts, experience SD-mediated CNS shutdown during environmental stress and in response to a variety of experimental treatments, including inhibition of the Na^+^/K^+^-ATPase (NKA) using ouabain (Andersen et al., 2022; Robertson et al., 2020; Robertson et al., 2017; Rodgers et al., 2010; Spong et al., 2016a). Conversion of the NKA from a pump into a channel using palytoxin (Rossini and Bigiani, 2011) in both mammalian (Kim et al., 2025) and insect (Wang et al., 2024) preparations generates SD that mimics the effect of anoxia, indicating a vulnerability of the pump and suggesting that similar molecular mechanisms could be involved. In locusts, ouabain also generates a slow reversal of the transperineurial potential (TPP) measured across the BBB (Robertson and Van Dusen, 2021; Van Dusen et al., 2020) and this is similar to the NUP (Robertson and Wang, 2025). Pretreatment with the glycolytic inhibitor, monosodium iodoacetate (MIA), causes increased levels of damage such as protein oxidation and cellular apoptosis in the CNS, and this is associated with incomplete recovery of the TPP after anoxic SD in locusts (Wang 王宇扬 et al., 2023). The same damaging MIA pretreatment also accelerates the ouabain-induced NUP (RM Robertson, unpublished observations).

Recently, we found that the ouabain-induced SD and NUP are separable into distinct processes in the locust CNS and that the NUP is exacerbated by electrode damage to the BBB (Robertson and Wang, 2025). We also found that mitigation of oxidative tissue damage using the membrane permeable antioxidant, N-acetylcysteine amide (NACA), slows the development of the NUP. Given that SD represents a collapse of ion gradients due to transient or sustained failure of ion homeostasis, and that NUPs represent accumulating tissue damage, somehow associated with the BBB in locusts, and possibly associated with the BBB in humans (Dreier et al., 2019), we investigated ion homeostasis of the locust CNS with a particular focus on the BBB.

The development, structure, and function of insect glial cells is best described for *Drosophila*, which is promoted as a model for mammalian glia, notably with respect to human health concerns (Contreras and Klämbt, 2023; Fernandes et al., 2024; Rey et al., 2023; Yildirim et al., 2019). There is considerable evidence for evolutionary conservation of molecular mechanisms of the BBB (Hindle and Bainton, 2014; Hindle et al., 2017; Stork et al., 2008). The sheath encasing the insect CNS comprises an extracellular lamella covering perineurial and subperineurial glial layers (Limmer et al., 2014) and the subperineurial layer provides the barrier to the free flow of ions, small compounds and xenobiotics from the haemolymph. Paracellular diffusion is prevented by tight or septate junctions between the cells of the layer whereas intercellular diffusion is enabled by gap junctions. Whereas local mechanisms exist within the neuropil to regulate ion homeostasis in the immediate extracellular environment of neural circuits, the BBB is concerned with ion homeostasis of the CNS as a whole, to protect the neuropil from the extreme variation of ion concentrations in the haemolymph.

Early electrophysiological investigations of insect CNS ion homeostasis focused on connectives in cockroaches due to the ability to record intracellularly from axons and demonstrate a barrier to ion diffusion across the BBB after greatly increasing the potassium ion concentration ([K^+^]) in the bathing medium (Schofield and Treherne, 1984; Treherne and Schofield, 1979; Treherne and Schofield, 1981). The properties of the BBB overlying the neural integrating centres, the ganglia, are markedly different from those of the BBB at the connectives. High [K^+^] in the bathing solution generates repetitive SD measured by recording the TPP at a ganglion but not at connectives (Robertson et al., 2020; Schofield, 1990). Net current flow out of ganglia is associated with a high density of mitochondria in perineurial cells at those locations (Smith and Shipley, 1990) and the scanning ion electrode technique (SIET) has shown that this current is carried by K^+^ ions (Kocmarek and O’Donnell, 2011).

SD is associated with a surge of extracellular potassium ion concentration in the interstitium ([K^+^]_o_) and can be triggered by somatosensory stimulation in mice (von Bornstädt et al., 2015) and by activity-dependent increases of [K^+^]_o_ in locusts (Spong et al., 2016b). SD can cause profound cell swelling (Kirov et al., 2020), which can be monitored by an increase in the resistivity of the tissue (Traynelis and Dingledine, 1989; Walch and Fiacco, 2022). At locust ganglia the resistance to current flow from an extracellular electrode in the neuropil to the bathing solution increases during anoxic SD, which could be due to cell swelling and/or ion conductance changes at the BBB (Robertson and Van Dusen, 2021). Manipulating extracellular osmolarity to cause cell swelling (or shrinkage) exacerbates (or mitigates) the severity of ouabain-induced SD (Spong et al., 2015). Moreover, impairing the diffusion barrier function of the BBB with urea (Spong et al., 2014) or inhibiting gap junctions with carbenoxolone (Spong and Robertson, 2013) can trigger repetitive SD in the absence of any other treatment (e.g. ouabain or high [K^+^]).

Cation-chloride cotransporters play important roles in the regulation of cell volume of both insects (Leiserson et al., 2011; Leiserson and Keshishian, 2011) and mammals (MacAulay, 2021; Steffensen et al., 2015). Inhibition of the Na^+^/K^+^/2Cl^−^cotransporter (NKCC) using bumetanide exacerbates ouabain-induced SD (Spong et al., 2015) and eradicates a protective effect of rapid cold hardening that is associated with increased levels of NKCC protein in the CNS (Srithiphaphirom et al., 2023). The K^+^/Cl^-^cotransporter (KCC) also has a major role in cell volume regulation (Russell, 2000; Walch and Fiacco, 2022) and may be involved in SD-induced dendritic beading in mice (Steffensen et al., 2015) but its participation in locust SD is unknown. The possibility that KCC could be involved in regulating insect SD is indicated by the observations that KCC is enriched in subperineurial glia of *Drosophila*, and glial-specific knockdown of the KCC gene, *kazachoc*, causes cell swelling and promotes a seizure-sensitivity in the CNS that is associated with a vulnerability to chill coma (= SD) (Hekmat-Scafe et al., 2006; Rusan et al., 2014).

We investigated mechanisms of ion homeostasis of the locust CNS to test the hypothesis that the ionoregulatory role of the BBB protects against SD and the NUP induced by ouabain. Insects can be exposed to cardiac glycosides like ouabain in their diet and the perineurium of the CNS is an effective diffusion barrier to these compounds (Petschenka et al., 2012; Petschenka et al., 2013). In *Drosophila*, an active transport barrier for ouabain co-located with NKA has been described (Torrie et al., 2004). In addition, although sensitivity to ouabain is conserved across taxa (Blaustein and Hamlyn, 2024), amino acid substitutions near the ouabain-binding pocket of the NKA in *L. migratoria* may reduce its sensitivity to ouabain (Dobler et al., 2015; Yang et al., 2019). Simultaneous intracellular recording from neuronal dendrites and extracellular recording of the TPP was used to assess the contribution of neuronal activity to ouabain vulnerability. We used SIET to measure K^+^ flux across the BBB at ganglia and the connectives, before and after sodium azide-induced SD. The subperineurial layer (= BBB) of the *Drosophila* CNS contains an active efflux transporter (an ATP binding cassette transporter, Mdr65) that is protective against cytotoxic xenobiotics, homologous to the mammalian transporter MDR1/Pgp, and inhibited by cyclosporin A (Mayer et al., 2009). This is also true of the locust BBB (Andersson et al., 2014). We compared the impact on ouabain-induced SD and NUP of impairing the diffusion barrier (by cutting the nerve roots) with impairing the transport barrier (by pretreatment with cyclosporin A) and with impairing both together. To inhibit the KCC we pretreated with 4,4ʹ-diisothiocyanostilbene-2,2ʹ-disulfonic acid (DIDS) (Delpire and Lauf, 1992; Russell, 2000).

## Materials and Methods

### Animals

A breeding colony of migratory locusts (*Locusta migratoria*) was maintained in the Department of Biology at Queen’s University, Kingston, Canada. Cages were crowded to ensure the locusts remained in the gregarious phase and they were illuminated with 40 W incandescent light bulbs during a 12-hour light period, which raised cage temperatures to ∼30 °C from ∼26 °C during the dark period. The locusts were fed daily with wheat grass and a dry mixture of yeast, bran, and milk powder *ad libitum*. Mature (3 − 7 weeks since the imaginal moult) male and female locusts were randomly assigned to treatment groups for different experiments, and treatments were not blinded during data collection or analysis. Group sizes and sex composition for experiments are described in the Results section. For experiments comparing different sexes, preparations or treatments, locusts from the same cage were used each day of experiments to control for colony conditions and the order of experiments was arranged to control for time of day.

### Electrophysiological Recording

Semi-intact preparations (Robertson and Pearson, 1982) exposed the thoracic ganglia for intracellular and extracellular recording. The legs and wings were removed prior to pinning a locust to a cork platform, and the thorax was opened with a dorsal midline incision. The gut was reflected out of the cavity and the ganglia were revealed by removing overlying tissues (salivary glands, gonads and musculature). To stabilize the ganglia for intracellular recording, nerve roots 3, 4 and 5 on both sides of the meso- and metathoracic ganglia were cut approximately ¼ mm from the ganglia, which were then lifted onto a stainless-steel plate. For TPP recording with an extracellular electrode, the ganglia were not lifted onto the plate and the nerve roots were cut or not, depending on the experiment. Preparations were grounded using a chlorided silver wire placed in the abdomen and bathed in standard locust saline (in mM: 147 NaCl, 10 KCl, 4 CaCl_2_, 3 NaOH, 10 HEPES buffer; pH adjusted to 7.2; chemicals from Sigma-Aldrich).

Intracellular electrodes were pulled from borosilicate glass capillaries (WPI Inc, Sarasota, FL) using a P-97 Flaming/Brown puller (Sutter Instrument Co., Novato, CA) to a resistance of 20-40 Megohm when back-filled with 500 mM KCl. Extracellular electrodes were similarly pulled to a resistance of 5-7 Megohm back-filled with 3 M KCl. Brief (200 ms) current pulses were delivered through extracellular and intracellular electrodes to measure the resistance across the BBB or across neuronal membranes, respectively. Signals were amplified using model 1600 Neuroprobe amplifiers (A-M systems Inc., Carlsborg, WA), digitized using a 1440 A digitizer and recorded using Axoscope 10.7 for later analysis with Clampfit 10.7 (Molecular Devices LLC, San Jose, CA). Digital sampling rate was 10 kHz for intracellular recording but reduced to 1 kHz if only the extracellular TPP was recorded.

### K^+^ flux measurement

K⁺ fluxes were measured from mesothoracic ganglia of *in vitro* preparations (see Fig. 3A), using adult female locusts, as their larger size facilitated a higher success rate in dissection. Prior to dissection the animals were transferred and housed in well-ventilated, clear acrylic enclosures (30 × 20 × 15 cm, 10 animals per cage). Isolation of the nerve cord was necessary to facilitate electrode placement and eliminate K⁺ efflux interference from muscles (RA Van Dusen and RM Robertson, unpublished observations). Tissue oxygenation was maintained by preserving the main tracheal reservoir and using aerated bathing saline. In a semi-intact preparation, the glands and muscles covering the thoracic ganglia were removed with forceps. Peripheral nerve roots were severed approximately 0.5 mm from the ganglia surfaces bilaterally. The main ventral tracheae were cut anterior to the head and posterior to the first abdominal ganglion, while the communicating tracheoles were severed, except for those supplying the ganglia. Finally, the connectives rostral to the prothoracic ganglion and caudal to the first abdominal ganglion were cut, and the isolated ganglia-tracheae assembly was immediately transferred into a Sylgard-lined glass petri dish (60 mm diameter) containing aerated standard locust saline. The ganglia were immobilized in the dish with the dorsal side facing up and pinned between the connectives to minimize further damage.

#### Electrode Construction

K^+^ selective microelectrodes were produced as previously described in detail (Donini and O’Donnell, 2005). Thin walled unfilamented glass capillaries (TW150-4, WPI, Sarasota, Fl, USA) were pulled on a P-97 Flaming/Brown puller (Sutter Instrument Co., Novato, CA) to a tip diameter of ∼ 5µm. Capillaries were silanized before backfilling with 100 mM KCl and then front filled with K^+^ ionophore I, cocktail B (Millipore Sigma, Oakville, ON. Canada). The reference electrodes were produced by filling glass capillaries (TW-150-4) with a simmering aqueous solution of 3 mol l-1 KCl containing 2% agar. Filled capillaries were cooled at room temperature to solidify the agar solution.

#### SIET Protocol

The lateral surfaces of the mesothoracic ganglion between N1 and N2, as well as the connective between meso- and metathoracic ganglia were selected for SIET measurements to ensure optimal electrode access. The SIET technique employed in this study has been described in detail elsewhere (Donini and O’Donnell, 2005). The K⁺ electrode was calibrated in 1 and 10 mM KCl, the slopes for 10-fold changes in [K⁺] were [mean ± S.D.] 58.3 ± 3.2 mV (N = 4). The electrode excursion distance was set to 200 µm, with wait and sample periods of 3 s and 1 s, respectively. Voltage gradients were recorded as the average of three repeated differential measurements using the sampling protocol described, starting at the ganglion surface.

Background noise levels measured using the same sampling protocol approximately 5 mm away from the tissue preparation were subtracted from the average voltage gradient measured at the preparation. The concentration gradient was then calculated using the voltage gradient using the following formula:

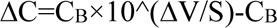

where ΔC represents the concentration gradient in μmol l^-1^ cm⁻³, C_B_ is the background [K^+^] concentration, calculated as the mean of the concentration at each point measured with the electrode in μmol l^-1^, ΔV is the mean voltage gradient recorded at the point of interest (after subtracting background), and S is the average slope of the electrode during calibration. K⁺ fluxes were calculated from the concentration gradient using Fick’s law of diffusion.

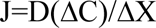

where J is the net flux of K^+^ (pmol cm^-2^ s^-1^), D is the diffusion coefficient of K^+^ 1.19 x 10^-5^ cm^-2^ s^-1^, ΔC is the concentration gradient (pmol cm^-3^) and ΔX is the distance between the two points measured (cm).

In preliminary experiments we found that low concentrations of ouabain (10⁻⁴ to 10⁻^2^ M) permanently impaired the K⁺ ionophore in the electrode, leading to unpredictable signal drift during calibration. Hence, we used sodium azide, which did not interfere with the ionophore, to generate sustained SD in the following procedure. All SIET and accompanying extracellular measurements were completed within 15 minutes of the initial extraction of the ganglia to ensure tissue viability. Following dissection, background noise in the bathing saline was measured in the Sylgard dish, followed by the measurement of pre-SD gradients at the ganglion surface and connective. The dish was then immediately transferred to the DC workstation, where an extracellular DC electrode measured TPP in the mesothoracic ganglion. The aerated saline was carefully drained with a pipette and replaced with 10 mM sodium azide in standard locust saline to mimic anoxic SD. Once the abrupt negative DC shift of TPP occurred, the dish was transferred back to the SIET workstation, where background noise was measured again in the presence of azide. The SIET electrode was then visually repositioned near the pre-SD measuring site, before the post-SD gradients near the ganglion surface and connectives were measured.

### Pharmacology

Ouabain, 4,4ʹ-diisothiocyanostilbene-2,2ʹ-disulfonic acid (DIDS) and cyclosporin A were purchased from Sigma-Aldrich. Cyclosporin A and DIDS were made up as stock solutions of 1 mM in locust saline and frozen in aliquots prior to use. For both pre-treatments, saline was removed from the thoracic cavity and the interior was dried with a paper tissue before treatment. For cyclosporin A, 1 mL of 1 mM was added to the preparation (e.g. (Andersson et al., 2014)) and, depending on the experiment, nerves 3, 4 and 5 of the meso and metathoracic ganglia were cut, which improved access to the neuropil. For DIDS, 0.5 mL of saline was returned to the preparation and 0.5 mL of 1 mM DIDS was added to achieve a final concentration of 500 µM (e.g. for similar effective concentrations in insects (Schewe et al., 2012) and (Del Duca et al., 2011)). After 20 minutes pre-treatment, 1 mL of 10 mM ouabain was added to achieve a final concentration of 5 mM of ouabain, which is an effective procedure for induction of TPP reversal and SD in preparations with cut nerve roots (Robertson and Van Dusen, 2021) (Robertson and Wang, 2025). TPP was recorded for at least 30 minutes. If ouabain failed to generate SD, mostly for preparations with intact nerve roots and without pretreatment, the preparation chamber was filled with nitrogen gas (see (Robertson and Van Dusen, 2021)) to generate anoxic SD and confirm the reliability of the TPP recording.

### Data Analysis

Ouabain treatment of the locust CNS generates repetitive SD events in neuropil superimposed upon a slow reversal of the TPP from positive to negative, which resembles the NUP of mammalian CNS (Robertson and Wang, 2025; Van Dusen et al., 2020; Wang et al., 2024) (**Fig. 1A**). We used Clampfit 10.7 to analyse the traces. When necessary, they were filtered to remove high frequency electrical noise and/or 60 Hz AC interference. To characterize the rate and extent of ouabain-induced NUP and its effect on the propensity to generate SD events, we measured: 1) initial TPP, 2) early positive shift induced by ouabain, 3) TPP 30 minutes after the onset of ouabain, 4) latency to the first SD event, 5) number of SD events during 30 minutes of ouabain treatment, and 6) SD mean frequency calculated from the average period between all SD events in all preparations. The last two measures are related but differ in that SD frequency is measured for events occurring after the latent period. For statistical analysis, preparations which did not generate an SD event within 30 minutes were assigned a latency = 31 mins, number of SDs = 0, and frequency = 0 /10 mins. To characterize the trajectory of the TPP during SD events, we measured: i) maximum SD amplitude, ii) duration measured at half amplitude, iii) slope of TPP at SD onset, iv) slope of TPP recovery, v) time to recover taken from the negative peak to half amplitude of recovery, and vi) the time constant (tau) of an exponential curve fitted to the TPP recovery trajectory using Clampfit. To perform statistical tests and generate graphs we used Sigmaplot 15 (Systat, Grafiti LLC, Palo Alto, CA) and GraphPad Prism 10 (GraphPad Software, Boston, MA). Outliers were identified using Grubb’s Test online (GraphPad) and removed from further analysis. Data were tested for normality (Shapiro-Wilk Test) and equal variance (Browne-Forsyth Test) and statistical tests were applied as described in the Results. Some of the data did not satisfy the normality and equal variance requirements for parametric ANOVAs. Nevertheless, we used parametric 3-way and 2-way ANOVAs to analyse the data. With respect to type 1 error, the F-test has 100% robustness for violations of normality showing that parametric ANOVAs are valid procedures for data that are not normally distributed (Blanca et al., 2017). In addition, for the combination of variance ratios and group size ratios of our data, the F-test for type 1 error has 100% robustness for violations of equal variance (Blanca et al., 2018). This allowed us to use the more powerful parametric ANOVAs and enabled multiple pair-wise comparisons within sex, preparations and treatments. Nonparametric Kruskal-Wallis 1-way ANOVA on ranks applied independently to analyse the effects of sex, preparation or treatment confirmed the validity of using parametric ANOVAs by giving almost identical P values (not shown). Descriptive results in the text are given as mean ± standard deviation (SD). Data for comparison are presented in bar graphs showing mean ± standard error of the mean (SEM) overlaid with individual data points.

**Figure 1.**
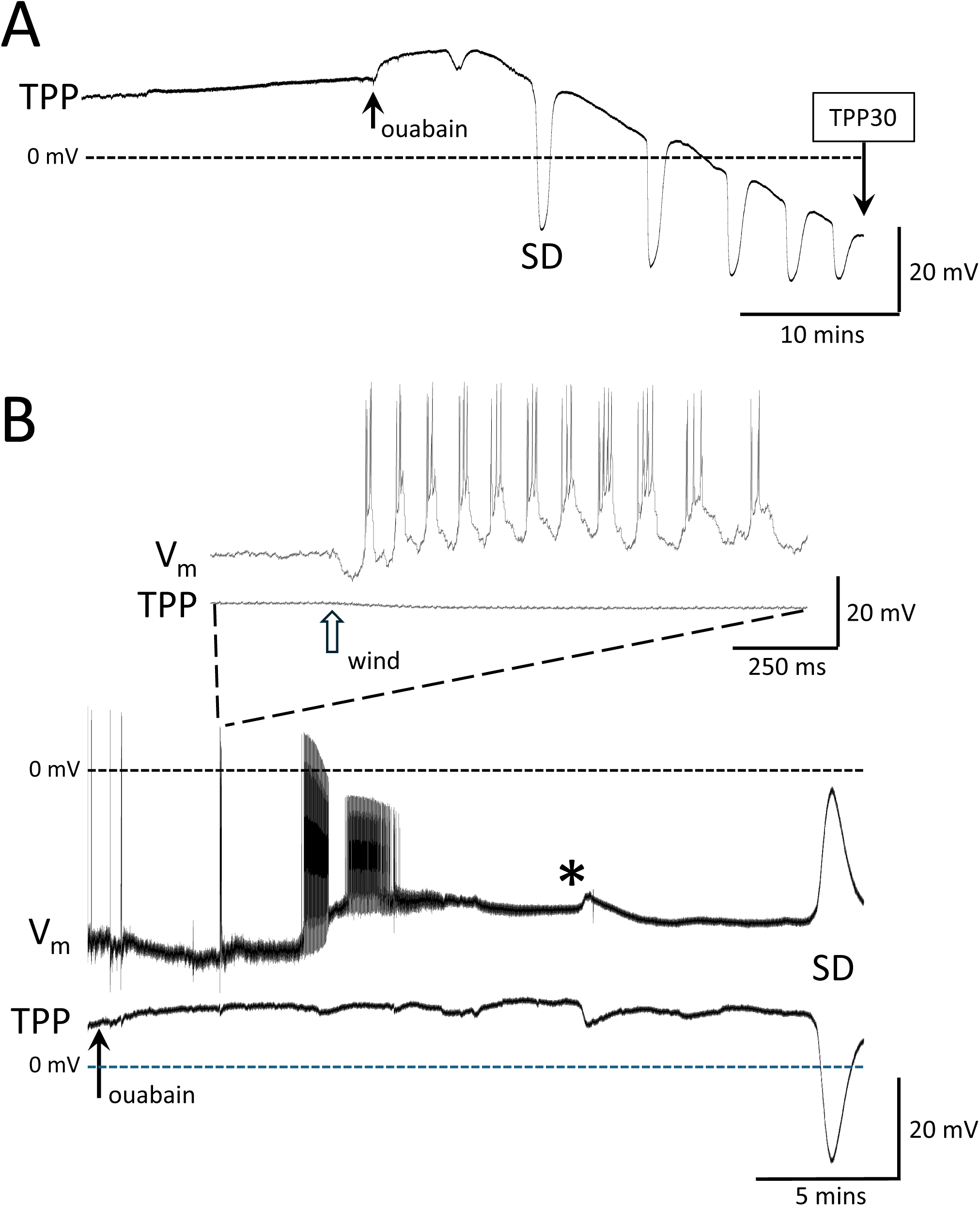
Ouabain induces spreading depolarization (SD) and a slow reversal of the transperineurial potential (TPP) from positive to negative. A. Recording of TPP from a metathoracic ganglion responding to treatment with ouabain (arrow). Note that ouabain induced a positive shift followed by a reversal from positive to negative superimposed with SD events. The rate of TPP reversal is characterised by the potential reached 30 minutes after the onset of ouabain treatment (TPP30). B. Intracellular recording of a forewing depressor motoneuron in the mesothoracic ganglion after the application of ouabain (arrow) at the beginning of the trace. The expanded portion (top) illustrates generation of a brief flight motor rhythm in response to a wind stimulus (open arrow) to the head of the preparation. Note that the thickness of the V_m_ trace during the first 10 minutes is due to a continuous barrage of synaptic potentials. The asterisk indicates the time at which synaptic potentials could no longer be identified in the neuronal trace. TPP − transperineurial potential;V_m_ − neuronal membrane potential.

**Figure 2.**
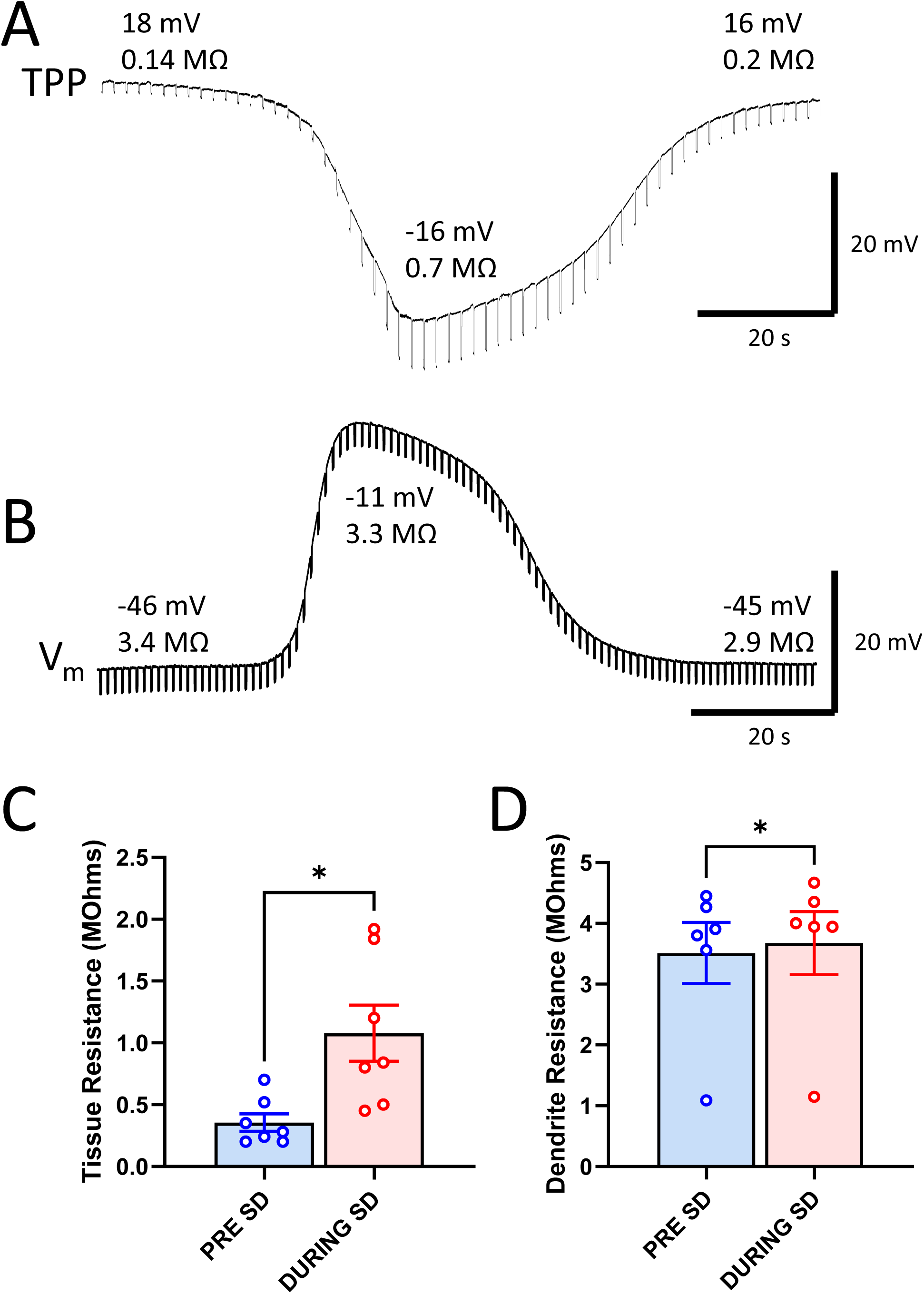
Ouabain-induced SD is associated with an increase in resistance across the BBB but negligible change in neuronal membrane resistance. A. Extracellular recording of the TPP of the metathoracic ganglion during SD in the presence of ouabain. 10 nA, 200 ms current pulses were delivered at 0.5 Hz to measure the resistance to current flow across the BBB. **B.** Intracellular recording of the membrane potential (V_m_) from the dendrite of a forewing depressor motoneuron in the mesothoracic ganglion. 1 nA, 200 ms current pulses were delivered at 1 Hz to measure input resistance of the dendrite. In both **A** and **B**, the recorded potential (mV) and the resistance (MΩ) is noted before, during and after the SD event. **C.** Tissue resistance increased approximately three-fold during SD. **D.** Input resistance of the motoneuron dendrite increased a small amount during SD. Graphs show mean ± SEM overlaid with individual data points. *- P < 0.05

**Figure 3.**
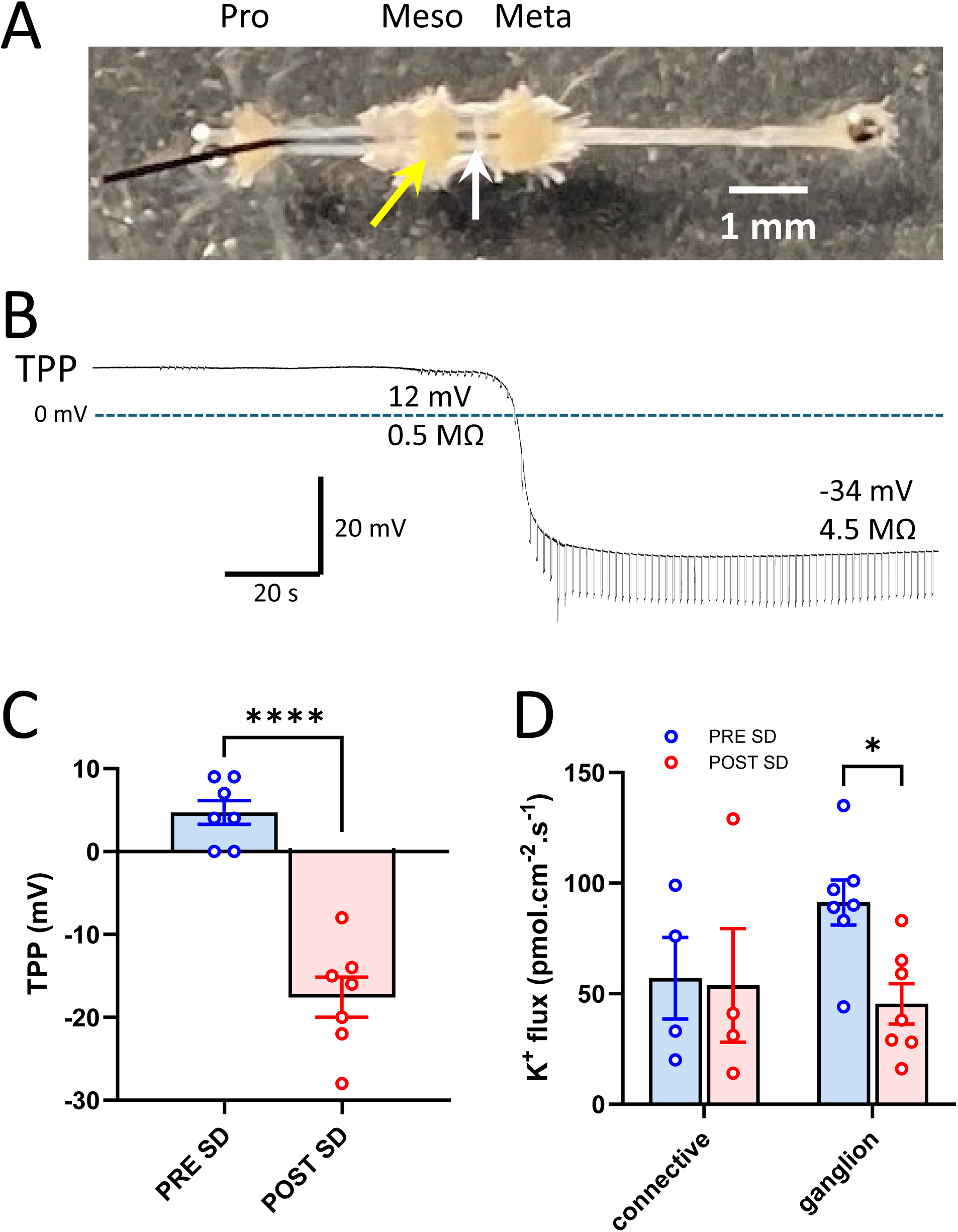
Increased resistance across the BBB is associated with a decrease in K^+^ efflux from the ganglion but not from the connective. A. Isolated preparation of the pro-, meso-, and metathoracic ganglia indicates location of K^+^ electrode for measuring flux at the mesothoracic ganglion (yellow arrow) or at the meso-meta connective (white arrow). **B.** Recording of the TPP of the metathoracic ganglion in an isolated preparation. The trace starts 45 minutes after the electrode was inserted into the ganglion and 5 minutes after the addition of 10^-2^ M ouabain to the dish. 2.5 nA, 200 ms current pulses were delivered at 1 Hz to measure resistance across the BBB. The recorded TPP and the resistance is noted before and after the occurrence of a sustained SD. **C.** TPP measured from the mesothoracic ganglion in isolated preparations before (PRE SD) and after (POST SD) the occurrence of SD in response to treatment with 10 mM sodium azide confirms that sustained SD was generated. **D.** In the same preparations, measurement of K^+^ efflux from the meso-meta connective was unaffected by SD, whereas K^+^ efflux from the mesothoracic ganglion was halved after SD. Graphs show mean ± SEM overlaid with individual data points. *- P < 0.05; **** - P < 0.0001.

## Results

### Neuronal responses

We recorded intracellularly from the dendrites of 17 wing muscle motoneurons in meso- and metathoracic ganglia (11 males, 6 females), identified by their bursting activity in response to a wind stimulus to the head (**Fig. 1B**). There were no statistical differences in any measure due to sex, so the data were pooled. Neuronal membrane potential (V_m_) before ouabain application was -57 ± 10 mV (mean ± SD). Ouabain application had a relatively rapid effect, depolarizing neurons by 17 ± 7 mV and generating a burst of action potentials culminating in spike failure in 3.2 ± 0.9 mins. The first SD event occurred 9 ± 5.2 min after ouabain was applied, 6 ± 4.3 mins after the neurons stopped firing. TPP recorded simultaneously started at 15 ± 6.6 mV and the amplitude of the SD event was 32 ± 5.4 mV.

Current pulses across the BBB, delivered via the extracellular TPP recording electrode in 7 preparations, revealed that the tissue resistance increased approximately three-fold from 0.4 ± 0.06 MΩ (mean ± SEM) prior to SD to 1.1 ± 0.20 MΩ at the negative peak of the first SD event (Paired t-test, P = 0.012) (**Fig. 2A,C**). However, current pulses delivered in 6 intracellular recordings showed only a small increase in dendrite membrane input resistance during SD (pre-SD: 3.5 ± 0.5 MΩ; during SD: 3.7 ± 0.5 MΩ; Paired t-test, P = 0.046) (**Fig. 2B,D**).

Summary: Flight motoneurons depolarized and lost excitability relatively rapidly after ouabain application. This occurred several minutes before the first SD event, which subsequently depolarized Vm close to 0 mV but was not associated with a decrease in neuronal membrane resistance as has been documented for flight motoneurons experiencing anoxia-induced SD (Robertson and Van Dusen, 2021).

### K+ efflux across the BBB

To determine the effect of SD on BBB activity at the hemolymph interface, we measured K⁺ fluxes from the surface of isolated mesothoracic ganglia and connectives (**Fig. 3A**) using SIET. In preliminary experiments, TPP measurement of isolated ganglia after application of ouabain confirmed the viability of the *in vitro* preparation for this experiment by demonstrating an abrupt negative shift in TPP associated with an increase in BBB resistance (**Fig. 3B**). For experiments measuring K^+^ flux before and after SD, SD was induced by an azide bath, following the initial flux measurement, and was confirmed by a negative shift of TPP (**Fig. 3C**, mean ± SEM, pre-SD TPP: 4.7 ± 1.4 mV, post-SD TPP: -17.6 ± 2.4 mV; n = 7, Paired t-test p < 0.0001). K⁺ efflux was detected at the surface of both the ganglia and connectives (pre-SD fluxes in pmol cm⁻² s⁻¹; ganglia: 91 ± 13, n = 7; connectives: 57 ± 16, n = 4). SD attenuated K⁺ efflux from the ganglion surface by 50%, while sparing the connectives (post-SD fluxes in pmol cm⁻² s⁻¹; ganglia: 45 ± 24, n = 7; connectives: 53 ± 51, n = 4). SD had a significant effect on K^+^ efflux from ganglia (Paired t-test p = 0.012) but not on K^+^ efflux from connective (Paired t-test p = 0.85) (**Fig. 3D**). The occurrence of SD reduced the initially greater ganglionic K⁺ fluxes to a level similar to that of the connectives.

Summary: During SD the increased resistance to current flow from the neuropil to the hemolymph was associated with a large reduction of the efflux of K^+^ ions from the surface of the mesothoracic ganglion while K^+^ flux from the connective was unaffected.

### Cyclosporin A pre-treatment

In 10 of 11 preparations with intact nerve roots and in control saline, no NUP or SD were observed within 30 minutes and we used nitrogen gas to induce anoxic SD and confirm the reliability of the recording (**Fig. 4A**). Impairing the BBB by cutting the nerve roots and/or pretreating with cyclosporin A accelerated the NUP and induced more SD events sooner (**Fig. 4B**). We investigated the effect of cyclosporin A pre-treatment on ouabain-induced SD in 51 successful preparations: 6 male control with intact nerve roots; 5 female control with intact nerve roots; 9 male control with cut nerve roots; 9 female control with cut nerve roots; 6 male cyclosporin A with intact nerve roots; 6 female cyclosporin A with intact nerve roots; 5 male cyclosporin A with cut nerve roots; 5 female cyclosporin A with cut nerve roots. Three-way ANOVAs revealed no effect of sex for any of the parameters we measured. Thus, the following description is for analysis with the sexes pooled.

**Figure 4.**
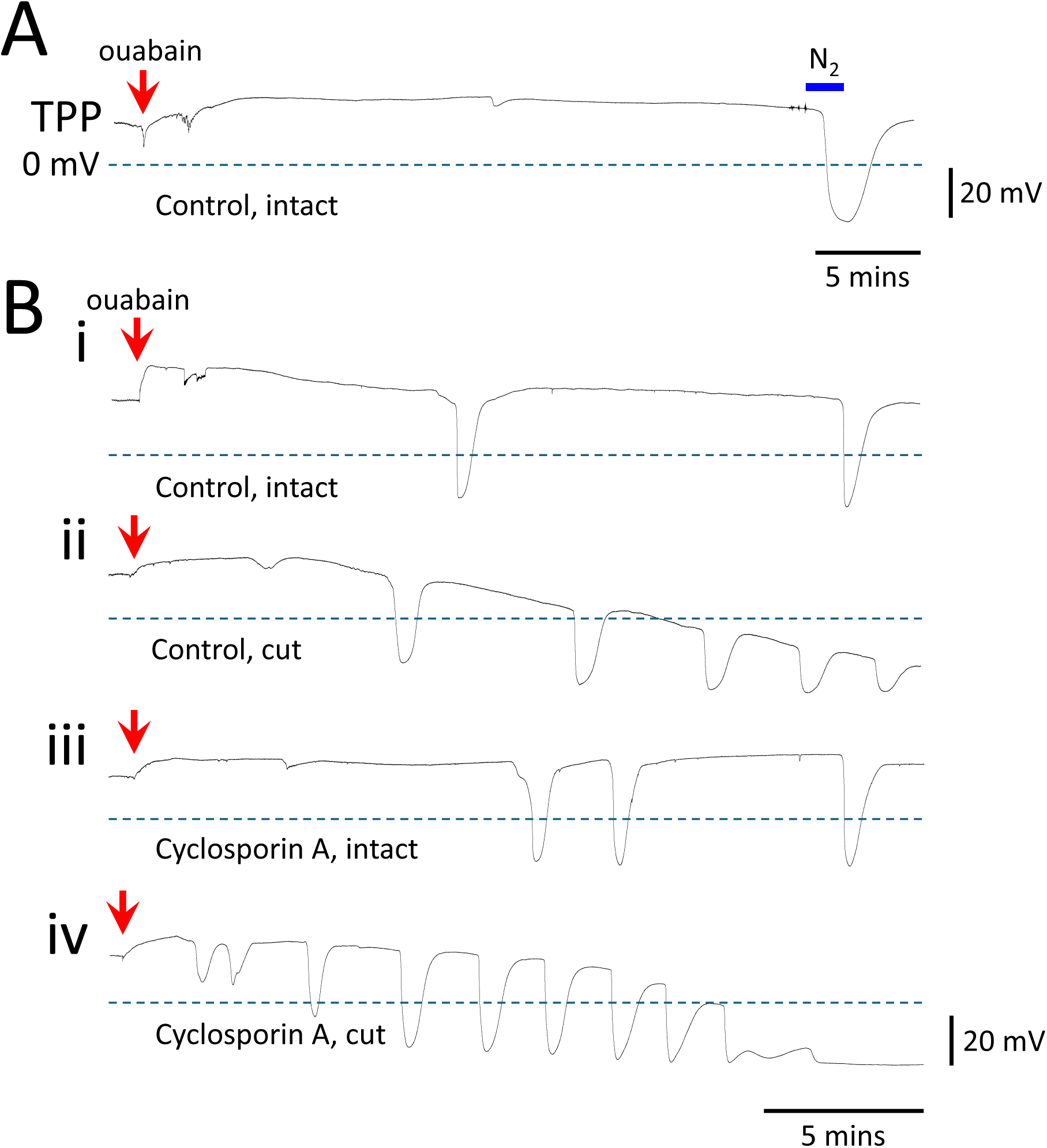
Impairment of the BBB exacerbates the NUP and promotes SD. A. Sample recording of TPP from a preparation with intact nerve roots and in control saline. After 30 mins with no SD events, an anoxic SD was induced with nitrogen gas (N_2_). **B.** Sample recordings from preparations: **i.** control saline and intact nerve roots (only one of eleven such preparations showed NUP and SD); **ii.** Control saline with cut nerve roots; **iii.** pretreated with cyclosporin A and intact nerve roots; **iv.** pretreated with cyclosporin A with cut nerve roots.

Cutting the nerve roots decreased the initial TPP (**Fig. 5A**) (mean ± SEM: n = 23 intact: 20 ± 1.1 mV; n = 28 cut: 15 ± 0.9 mV; 2-way ANOVA: P_roots_ = 0.001, P_treatment_ = 0.12). However, although the interaction did not reach significance (P_roots x treatment_ = 0.09), this result is driven by an effect of cyclosporin A to make the TPP more positive in intact preparations (Holm-Sidak multiple comparisons: P_treatment_ within intact = 0.028; P_treatment_ within cut = 0.92; P_roots_ within control = 0.20; P_roots_ within cyclosporin A = 0.001). This conclusion is supported by the observation that cutting the nerve roots decreased the positive shift of TPP induced by ouabain (**Fig. 5B**) (n = 23 intact: 11.5 ± 1.1 mV; n = 28 cut: 7.9 ± 0.78 mV; 2-way ANOVA: P_roots_ = 0.003, P_treatment_ = 0.1, P_roots x treatment_ = 0.54). Pretreatment with cyclosporin A enhanced the effect on the positive shift of cutting the nerve roots (Holm-Sidak multiple comparisons: P_treatment_ within intact = 0.46; P_treatment_ within cut = 0.1; P_roots_ within control = 0.06; P_roots_ within cyclosporin A = 0.017).

**Figure 5.**
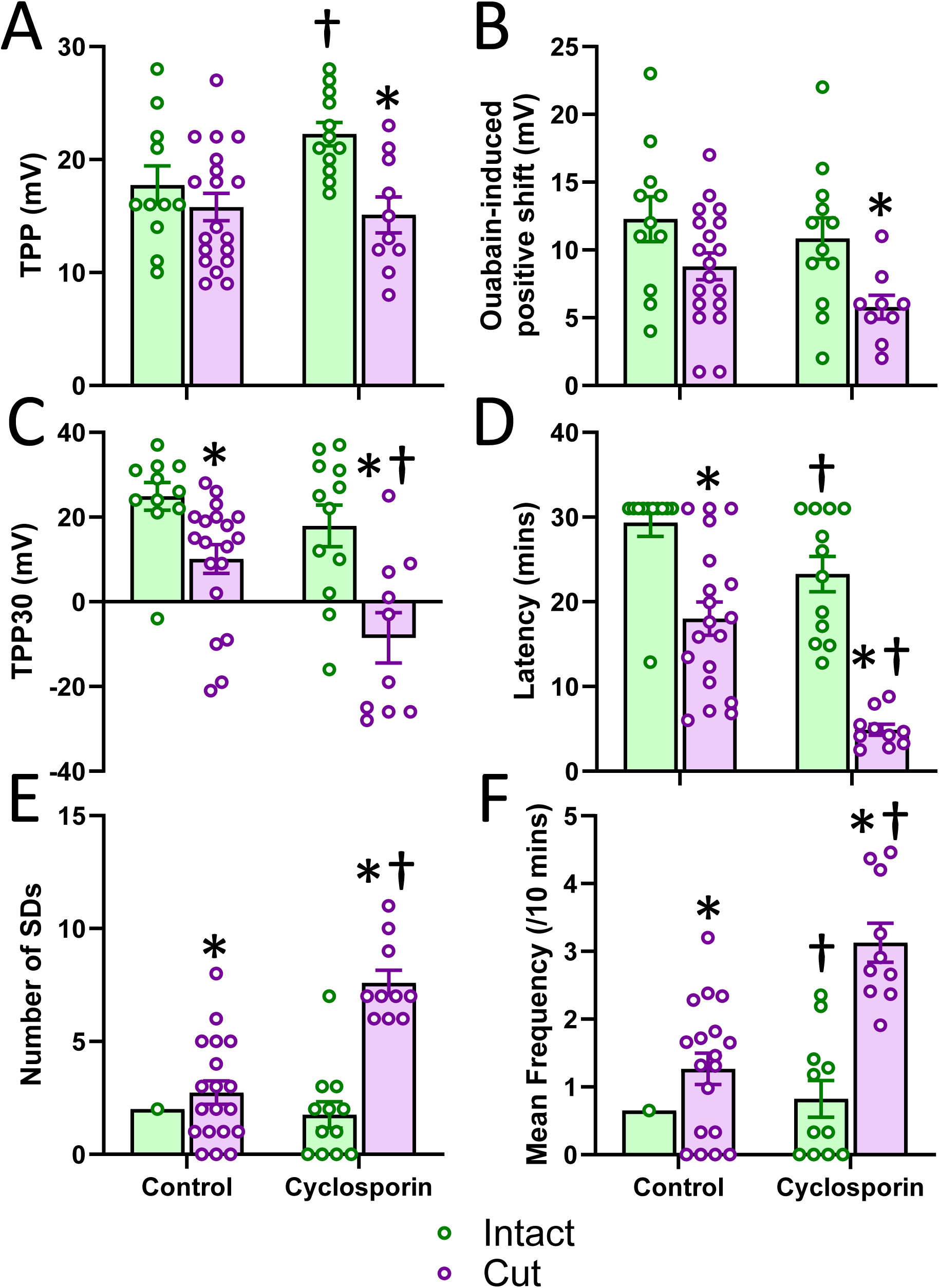
Cutting the nerve roots and/or treatment with Cyclosporin A exacerbates ouabain-induced NUP and evokes more SD events. A. Maximum amplitude of SD events. **B.** Duration of SD events. **C.** Slope of the TPP trajectory at SD onset. **D.** Slope of the TPP trajectory at SD recovery. **E.** Time to TPP recovery from the negative peak. **F.** Time constant (tau) of the exponential recovery of TPP at the end of SD events. Graphs show mean ± SEM overlaid with individual data points. * indicates a significant effect of preparation (intact or cut roots) within a treatment condition. † indicates a significant effect of Cyclosporin A within a preparation condition. Alpha = 0.05.

Cutting the nerve roots and cyclosporin A pre-treatment independently increased the rate of TPP reversal (**Fig. 5C**) (TPP30 for n = 23 intact: 22 ± 3.5 mV; n = 28 cut: 3.5 ± 3.5 mV; n = 29 control: 16.5 ± 2.8 mV; n = 22 cyclosporin A 5.9 ± 4.7 mV; 2-way ANOVA: P_roots_ < 0.001, P_treatment_ = 0.002, P_roots x treatment_ = 0.32). There was also an additive effect such that the combination of cyclosporin A and cut nerve roots had the greatest effect (Holm-Sidak multiple comparisons: P_treatment_ within intact = 0.13; P_treatment_ within cut = 0.003; P_roots_ within control = 0.005; P_roots_ within cyclosporin A < 0.001).

The results are similar for the latency to the first SD event. Cutting the nerve roots and cyclosporin A pre-treatment independently decreased the latency (**Fig. 5D**) (n = 23 intact: 26.2 ± 1.5 mins; n = 28 cut: 12.7 ± 1.7 mins; n = 29 control: 22.0 ± 1.8 mins; n = 22 cyclosporin A 14.9 ± 2.3 mins; 2-way ANOVA: P_roots_ < 0.001, P_treatment_ < 0.001, P_roots x treatment_ = 0.11). There was also an additive effect such that the combination of cyclosporin A and cut nerve roots had the greatest effect (Holm-Sidak multiple comparisons: P_treatment_within intact = 0.035; P_treatment_within cut < 0.001; P_roots_ within control < 0.001; P_roots_ within cyclosporin A < 0.001).

Not surprisingly, the results are similar for the number of SD events. Cutting the nerve roots and cyclosporin A pre-treatment independently increased the number of events (**Fig. 5E**) (n = 23 intact: 1.0 ± 0.35; n = 28 cut: 4.4 ± 0.60; n = 29 control: 1.7 ± 0.41; n = 22 cyclosporin A 4.4 ± 0.75; 2-way ANOVA: P_roots_ < 0.001, P_treatment_ < 0.001, P_roots_ x treatment = 0.003). The combination of cyclosporin A and cut nerve roots had the greatest effect (Holm-Sidak multiple comparisons: P_treatment_ within intact = 0.052; P_treatment_ within cut < 0.001; P_roots_ within control = 0.001; P_roots_ within cyclosporin A < 0.001).

Also not surprisingly, the results are similar for the frequency of SD events after the latent period. Cutting the nerve roots and cyclosporin A pre-treatment independently increased the frequency (**Fig. 5F**) (n = 23 intact: 0.44 ± 0.16 /10 mins; n = 28 cut: 1.9 ± 0.25 /10 mins; n = 29 control: 0.72 ± 0.16 /10 mins; n = 22 cyclosporin A 1.9 ± 0.31 /10 mins; 2-way ANOVA: P_roots_ < 0.001, P_treatment_ < 0.001, P_roots x treatment_ = 0.012). The combination of cyclosporin A and cut nerve roots had the greatest effect on SD frequency (Holm-Sidak multiple comparisons: P_treatment_ within intact = 0.028; P_treatment_ within cut < 0.001; P_roots_ within control < 0.001; P_roots_ within cyclosporin A < 0.001).

It was not possible to perform 3-way ANOVAs on the SD event parameters because the cell for female, intact, control was empty i.e. had no SDs to measure. Indeed, only one male, intact, control generated an SD within 30 mins. To determine if there were sex differences in this dataset, we performed t-tests (or Mann Whitney tests) to compare male and female data. Only the slope of TPP recovery (P = 0.024) and the time constant of TPP recovery (P = 0.018) were different in males and females, with males being slower to recover.

In sharp contrast to the effects on the NUP and the propensity to generate SD events, there were no effects of cutting nerve roots and/or pre-treatment with cyclosporin A on any of the parameters characterizing the TPP trajectory underlying individual SD events (**Fig. 6**). For the 38 preparations that generated at least one SD event, SD amplitude (**Fig. 6A**) was 39 ± 1.0 mV (2-way ANOVA: P_roots_ = 0.54, P_treatment_ = 0.94). SD duration (**Fig. 6B**) was 0.9 ± 0.05 mins (2-way ANOVA: P_roots_ = 0.31, P_treatment_ = 0.77). Onset slope (**Fig. 6C**) was -5.7 ± 0.77 mV/s (2-way ANOVA: P_roots_ = 0.78, P_treatment_ = 0.79). Slope of TPP recovery (**Fig. 6D**) was 0.82 ± 0.05 mV/s (2-way ANOVA: P_roots_ = 0.85, P_treatment_ = 0.99). Time to recovery (**Fig. 6E**) was 0.64 ± 0.04 mins (2-way ANOVA: P_roots_ = 0.49, P_treatment_ = 0.75). The time constant of TPP recovery (**Fig. 6F**) was 17.3 ± 1.1 s (2-way ANOVA: P_roots_ = 0.67, P_treatment_ = 0.94).

**Figure 6.**
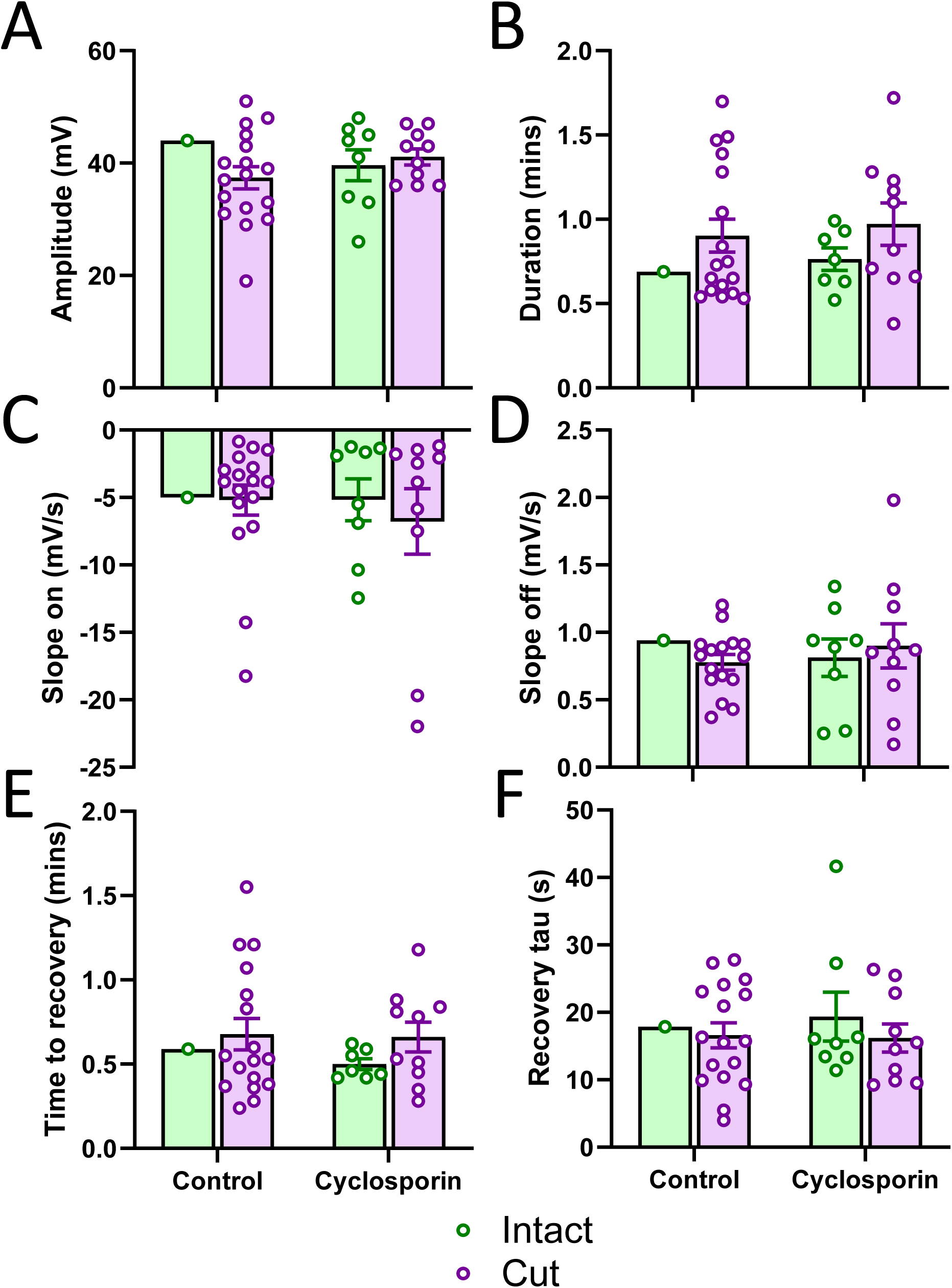
Cutting the nerve roots and/or treatment with Cyclosporin A does not affect parameters of the TPP trajectory of ouabain-induced SD events. A. Maximum amplitude of SD events. **B.** Duration of SD events. **C.** Slope of the TPP trajectory at SD onset. **D.** Slope of the TPP trajectory at SD recovery. **E.** Time to TPP recovery from the negative peak. **F.** Time constant (tau) of the exponential recovery of TPP at the end of SD events. Graphs show mean ± SEM overlaid with individual data points. Note that only one preparation (out of 11) with intact nerve roots and in the absence of drug treatment generated a ouabain-induced SD event within 30 mins.

Summary: Pre-treatment with cyclosporin A had an effect prior to the application of ouabain by making the TPP more positive. Both cutting the nerve roots to impair the BBB diffusion barrier and, independently, pre-treatment of intact ganglia with cyclosporin A to impair the BBB transport barrier strongly increased the rate of TPP reversal and promoted the generation of earlier, more frequent SD events. The effects of cutting the nerve roots and cyclosporin A were additive. The TPP recovery from SD events was faster in females.

### DIDS Pre-treatment

We investigated the effect of DIDS in 6 control males, 6 control females, 6 DIDS males and 6 DIDS females (**Fig. 7**). Following the pre-treatment, TPP (**Fig. 7A**) was 13.9 ± 1.24 mV (mean ± SEM; n = 24) and there was no effect of sex or treatment (2-Way ANOVA: P_sex_ = 0.95; P_treatment_ = 0.29). Application of ouabain induced a positive shift of 11.9 ± 0.92 mV (n = 24) (**Fig. 7B**) with no effect of sex or treatment (2-Way ANOVA: P_sex_ = 0.6; P_treatment_ = 0.2).

**Figure 7.**
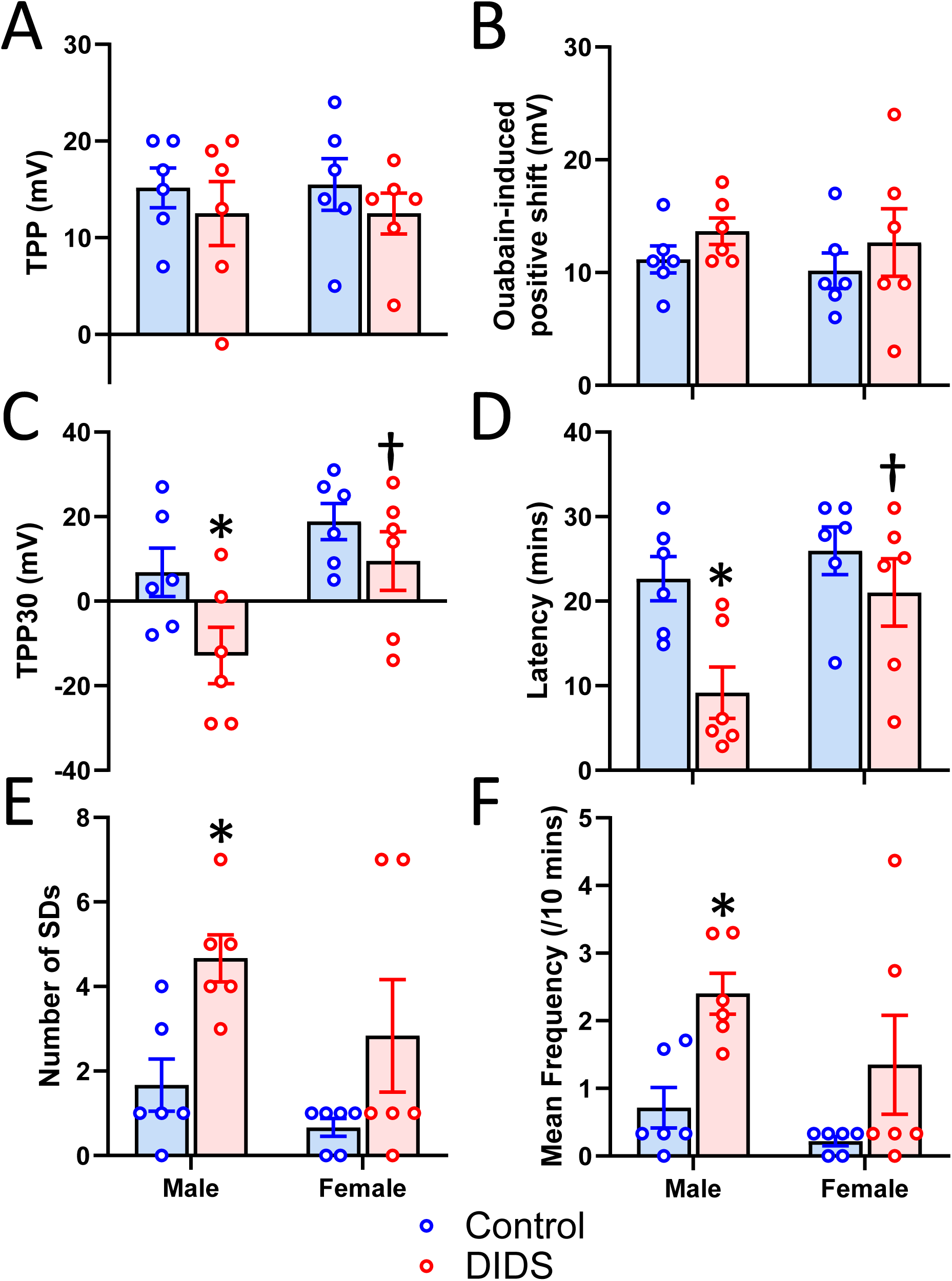
DIDS exacerbates ouabain-induced TPP reversal and evokes more SD events. Measurements of TPP recordings of ouabain-induced SD taken from male and female semi-intact preparations without (Control) or after treatment with 10^-3^ M DIDS. **A.** TPP prior to ouabain treatment. **B.** Positive shift of TPP after addition of ouabain. **C.** TPP at 30 minutes after addition of ouabain. **D.** Latency from ouabain addition to the first SD event. **E.** Number of SD events within 30 minutes after addition of ouabain. **F.** Mean frequency of SD events per 10 mins calculated from mean period between events. Graphs show mean ± SEM overlaid with individual data points. * indicates a significant effect of DIDS within a sex. † indicates a significant effect of sex within a treatment condition. Alpha = 0.05.

The rate of TPP reversal indicated by a negative TPP30 (**Fig. 7C**) was higher in males (in mV for n = 12; males: -3.0 ± 5.13; females: 14.2 ± 4.14) and after DIDS treatment (in mV for n = 12; Control: 12.8 ± 3.86; DIDS: -1.7 ± 5.69) (2-Way ANOVA: P_sex_ = 0.009; P_treatment_ = 0.025; P_sex x treatment_ = 0.39). The effect of DIDS was driven by a greater influence on males than females (Holm-Sidak multiple comparisons: P_treatment_ within males = 0.031; P_treatment_ within females = 0.28; P_sex_ within control = 0.17; P_sex_ within DIDS = 0.016).

The results are similar for the latency to the first SD event (**Fig. 7D**) which was shorter in males (in mins for n = 12; males: 15.9 ± 2.8; females: 23.5 ± 2.45) and after DIDS treatment (in mins for n = 12; Control: 24.3 ± 1.91; DIDS: 15.1 ± 2.98) (2-Way ANOVA: P_sex_ = 0.03; P_treatment_ = 0.009; P_sex x treatment_ = 0.19). The effect of DIDS was driven by a greater influence on males than females (Holm-Sidak multiple comparisons: P_treatment_ within males = 0.007; P_treatment_ within females = 0.28; P_sex_ within control = 0.47; P_sex_ within DIDS = 0.015).

The number of SD events (**Fig. 7E**) was not affected by sex (n = 12; males: 3.2 ± 0.6; females: 1.8 ± 0.72) but was increased by DIDS treatment (n = 12; Control: 1.2 ± 0.34; DIDS: 3.8 ± 0.74) (2-Way ANOVA: P_sex_ = 0.09; P_treatment_ = 0.004; P_sex x treatment_ = 0.6). The effect of DIDS was driven by an influence on males but not females, though the results for females are close to reaching significance (Holm-Sidak multiple comparisons: P_treatment_ within males = 0.014; P_treatment_ within females = 0.07; P_sex_ within control = 0.38; P_sex_ within DIDS = 0.12).

Not surprisingly, the results for frequency calculated from mean period of all SD events are similar (**Fig. 7F**). The frequency was not affected by sex (n = 12; males: 1.6 ± 0.33; females: 0.78 ± 0.39) but was increased by DIDS treatment (n = 12; Control: 0.47 ± 0.16; DIDS: 1.9 ± 0.41) (2-Way ANOVA: P_sex_ = 0.08; P_treatment_ = 0.003; P_sex x treatment_ = 0.52). The effect of DIDS was driven by an influence on males but not females (Holm-Sidak multiple comparisons: P_treatment_ within males = 0.011; P_treatment_ within females = 0.07; P_sex_ within control = 0.42; P_sex_ within DIDS = 0.09).

The maximum amplitude of SD events (**Fig. 8A**) was 35.7 ± 1.21 mV (n = 20) and was not influenced by sex or DIDS treatment (2-Way ANOVA: P_sex_ = 0.19; P_treatment_ = 0.82). Note that 4 preparations did not generate any SD events within 30 minutes. However, event duration (**Fig. 8B**) was longer in males (in mins n = 11 males: 1.1 ± 0.1; n = 9 females: 0.75 ± 0.05) but unaffected by DIDS treatment (in mins n = 9 Control: 0.88 ± 0.12; n = 11 DIDS: 0.99 ± 0.07) (2-Way ANOVA: P_sex_ = 0.02; P_treatment_ = 0.35; P_sex x treatment_ = 0.58) with no significant all pairwise comparisons.

**Figure 8:**
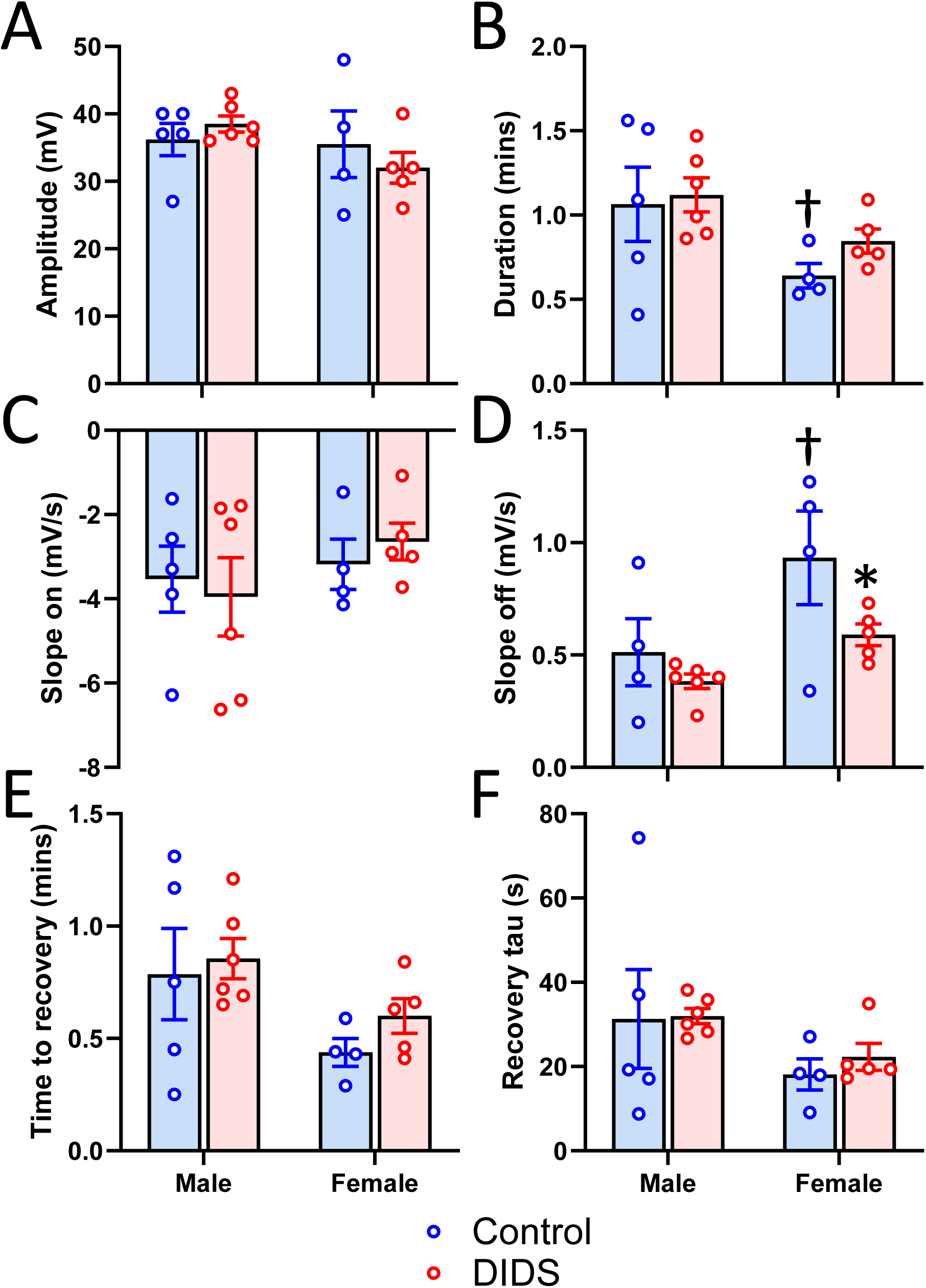
DIDS slows recovery of the TPP trajectory of ouabain-induced SD events in females. A. Maximum amplitude of SD events. **B.** Duration of SD events. **C.** Slope of the TPP trajectory at SD onset. **D.** Slope of the TPP trajectory at SD recovery. **E.** Time to TPP recovery from the negative peak. **F.** Time constant (tau) of the exponential recovery of TPP at the end of SD events. Graphs show mean ± SEM overlaid with individual data points. * indicates a significant effect of DIDS within a sex. † indicates a significant effect of sex within a treatment condition. Alpha = 0.05.

The slope of the TPP trajectory at SD onset (**Fig. 8C**) was -3.36 ± 0.34 mV/s (n = 20) and was not influenced by sex or DIDS treatment (2-Way ANOVA: P_sex_ = 0.3; P_treatment_ = 0.94). However, the slope of TPP recovery (**Fig. 8D**) was smaller in males (in mV/s n = 11 males: 0.59 ± 0.16; n = 9 females: 0.74 ± 0.09) but unaffected by DIDS treatment (in mV/s n = 9 Control: 0.88 ± 0.17; n = 11 DIDS: 0.48 ± 0.04) (2-Way ANOVA: P_sex_ = 0.003; P_treatment_ = 0.05; P_sex x treatment_ = 0.36). The effect of DIDS was driven by the influence of sex (Holm-Sidak multiple comparisons: P_treatment_ within males = 0.99; P_treatment_ within females = 0.03; P_sex_ within control = 0.005; P_sex_ within DIDS = 0.13).

The time to recovery of the TPP (**Fig. 8E**) was longer in males (in mins n = 11 males: 0.82 ± 0.09; n = 9 females: 0.53 ± 0.05) but unaffected by DIDS treatment (in mins n = 9 Control: 0.63 ± 0.11; n = 11 DIDS: 0.74 ± 0.07) (2-Way ANOVA: P_sex_ = 0.028; P_treatment_ = 0.37; P_sex x treatment_ = 0.71) with no significant all pairwise comparisons. The time constant of the TPP recovery (**Fig. 8F**) trajectory was 26.6 ± 2.89 s (n = 20) and not significantly affected by sex or treatment (2-Way ANOVA: P_sex_ = 0.09; P_treatment_ = 0.71).

Summary: Pretreatment with DIDS had a strong effect on the NUP induced by ouabain, speeding the rate of TPP reversal, which promoted the occurrence and increased the frequency of ouabain-induced SD events. This effect was more pronounced in males and multiple comparison tests revealed that the differences due to treatment reached significance only in males, though females showed similar trends. On the other hand, the TPP trajectory during SD events was marginally affected by DIDS. Females had shorter events, recovering more rapidly, and the only effect due to treatment was for the TPP recovery slope, a parameter which was significantly larger in females under control conditions.

## Discussion

Our primary goal was to test the hypothesis that ionoregulatory mechanisms of the locust BBB protect against SD and the NUP, which are electrical events associated with brain dysfunction and death in mammals. We found that treatments that impaired the diffusion and/or transport functions of the BBB accelerated the NUP leading to earlier and more frequent SD events. However, the same treatments had a negligible impact on the timing parameters of onset and recovery of SD, independent of the fact that females recovered ion homeostasis faster than males. Based on these and previously published results, we conclude that SD represents a transient, rapid, local failure of ion homeostasis in the neuropil, closely associated with neural cell swelling, whereas the NUP represents slowly accumulating oxidative damage to the perineurial cells of the BBB.

We also found that inhibition of NKA in the locust CNS caused a rapid, ∼17 mV depolarization of wing muscle motoneurons and silencing of neuronal activity, followed some time later by SD, which further depolarized the neurons to around 0 mV. During SD the resistivity of the tissue increased reversibly and there was a small increase (no decrease) in the input resistance of the neurons. K^+^ efflux from the neuropil (ganglion) but not the white matter (connectives), was substantially reduced during sustained SD. Repetitive SD was associated with a NUP.

Because they are easier to penetrate and identify based on their activity pattterns, our intracellular recordings were taken from the dendrites of wing muscle motoneurons and it would be dangerous to extrapolate the current findings to all neurons with dendrites or axons in the neuropil. Indeed, the structures and firing properties of flight motoneurons and flight interneurons, whose neurites are smaller in diameter, are quite different (Robertson and Pearson, 1982; Robertson and Pearson, 1983). Also the flight neuropil is generally confined to a dorsal layer just under the ganglion sheath (Ramirez and Pearson, 1988), suggesting that the flight circuitry may be the first to be affected by diffusion of ouabain into the ganglion in a semi-intact preparation supported by a ventral metal plate, and thus explain the lag between flight motoneuronal shutdown and SD. Nevertheless, neurons throughout the ganglion depolarize, fire a burst of action potentials and lose excitability in a continuous fashion whereas SD is an abrupt event, triggered when some threshold is crossed after all local excitability in the neuropil has shut down. Moreover, there was no indication of neuronal membrane conductance increase at onset or during SD that would indicate direct participation in generating SD. There was a slight increase in the measured neuronal input resistance with SD, but this is to be expected because the extracellular tissue is a part of the electrical circuit when passing current pulses through an intracellular electrode, and its resistivity increases several-fold. Previously we have described a neuronal conductance increase at the onset of SD (Robertson and Van Dusen, 2021) however in those experiments SD was triggered by anoxia, which would have had additional, direct effects on neurons, e.g., by impairing the energy supply. The notion that locust SD may be triggered by an abrupt failure of glial clearance mechanisms has been suggested for chilling-induced SD (Robertson et al., 2017) and is initiated close to the BBB (Robertson and Van Dusen, 2021) is supported by the results reported here.

Previous SIET measurements of K^+^ efflux across the BBB of a cockroach ganglion showed that metabolic inhibition with cyanide or NKA inhibition with ouabain increased K^+^ efflux from the ganglion (Kocmarek and O’Donnell, 2011). However, we found that metabolic inhibition with sodium azide to generate sustained SD markedly decreased K^+^ efflux from a ganglion. Moreover, given the close similarity between anoxic SD and ouabain-induced SD (Robertson and Wang, 2025), we would predict from this that SD induced by ouabain would also be accompanied by a reduction in K^+^ efflux from the interstitium to the hemolymph. The differences between our results are hard to reconcile. It may be due to a species difference (*P. americana* versus *L. migratoria*) but there are other, more consequential, technical differences.

We found that ouabain interfered with the K^+^ electrode, rendering the measurements of K^+^ flux in the presence of ouabain untrustworthy, and so we used sodium azide to generate SD. We have successfully used K^+^ electrodes and ouabain together in the past (e.g. (Rodgers-Garlick et al., 2011; Rodgers et al., 2007)) but in those instances the K^+^ electrode was implanted in a ganglion, through the protective sheath, alongside a reference electrode before ouabain was added to the bathing solution. Thus, the ionophore in the electrode tip was protected from the ouabain, which is not the case in the SIET studies. We also found that the ganglia were vulnerable to hypoxia during the procedure to isolate the nervous system in vitro, and without aeration/oxygenation of the saline and preservation of the tracheae, SD could occur during the setup, preventing measurement of K^+^ flux before SD (i.e., the initial TPP measurement was already substantially negative). Additionally, artifactual injury currents are generated and can interfere with measurements if the tracheae are cut close to the ganglion (Smith and Shipley, 1990). Finally, comparison with the previous SIET results is impossible without knowing the status of the cockroach preparation with respect to SD in the ganglion. We are confident that SD in our experiments was associated with a substantial decrease of K^+^ efflux from the ganglion paired with a marked increase in resistivity of the tissue. We suggest that cell swelling, which promotes SD (Spong et al., 2015), or more importantly, substantial shrinkage of an already restricted extracellular space (in a detailed ultrastructural study of the neuropil of the locust metathoracic ganglion, the extracellular distance between adjacent cell membranes is nowhere greater than 10 nm (Hoyle, 1986)), impedes the clearance of K^+^ ions from the interstitium to the hemolymph.

There was a sex difference in the response to ouabain after pretreatment with DIDS or cyclosporin A. A sex difference in the speed of recovery from anoxic SD (males slower) attributed to an increased metabolic rate in males has been noted and discussed previously (Robertson and Van Dusen, 2021). The same phenomenon is confirmed here for recovery from ouabain-induced SD events. In addition, we found a strong effect of sex on the rate of TPP reversal during the NUP after pretreatment with DIDS, but not after pretreatment with cyclosporin A, suggesting that males had increased vulnerability to damage. Females showed the same trends, which failed to reach significance. Such an effect could also be due to increased metabolic rate in males and increased damage from reactive oxygen species generated under stress. This proposition is supported by the fact that aging slows recovery from SD whereas pretreatment with the antioxidant NACA accelerated recovery from SD (Robertson and Wang, 2025), and in mammalian preparations the reduced speed of recovery with aging has been attributed to oxidative damage to NKA (Hertelendy et al., 2019).

We used cyclosporin A because it is established as an inhibitor of chemoprotective ABC transporters located in the BBB of *Drosophila* (Hindle et al., 2017; Mayer et al., 2009) and locusts (Andersson et al., 2014) but we are aware that it is commonly used as an immunosuppressant, prevents opening of the mitochondrial permeability pore (Hawrysh and Buck, 2013) and has also been shown to inhibit NKA (Burat et al., 2019). Similarly, we used DIDS with the intention of inhibiting the KCC (Delpire and Lauf, 1992; Russell, 2000) but recognize that it is a broad spectrum anion transport inhibitor that can target other ion homeostatic mechanisms involved in the control of cell swelling such as the sodium-activated chloride bicarbonate anion exchanger and chloride channels (Del Duca et al., 2011; MacAulay, 2021; Steffensen et al., 2015). Another concern is that bath applications of these agents will, over time, affect mechanisms in an increasing number of cell types in the CNS as they penetrate deeper into the tissue. Nevertheless, it is undeniable that cells of the BBB will be the first exposed to the inhibitors and thus are primary contenders to be responsible for the effects we describe here.

Pretreatment with DIDS accelerated the NUP and generated more SD events sooner. However, DIDS had minimal effect on the timing of the TPP trajectory during SD. Any significant DIDS pretreatment effects on individual SD events can be attributed to the strong effect of sex on recovery noted above. The results are clearer with cyclosporin A pretreatment such that it had strong effect on the NUP, measured by TPP30 and propensity to generate SD, and no effect on the parameters of SD. Damaging the diffusion barrier of the ganglion by cutting the nerve roots had similar consequences to pretreatment with cyclosporin A with a strong effect on NUP and no effect on SD. The effects of cyclosporin A pretreatment and cutting the nerve roots were additive. It was striking that exposure of intact nervous systems to ouabain for 30 mins was unable to generate SD events or a NUP in any females and in all but one male.

Cyclosporin A caused a positive shift of the initial TPP of intact nervous systems but had no impact on the relatively rapid positive shift of TPP caused by ouabain, which we attribute to ouabain inhibiting NKA in the basolateral membrane. We take this as an indication that cyclosporin A was not inhibiting NKA in our experiments. Previously we found that aging reduced the speed of recovery from SD and attributed this to age-related damage to NKA (Robertson and Wang, 2025). Thus, it is notable that while impairment of the BBB had strong effects on the propensity to generate SD and a NUP, there was no effect on the recovery from SD.

Based on the current and previous results, our updated conceptual model for SD and the NUP in insects is as follows. The mechanism for triggering SD is independent of the experimental method of inducing SD but different methods have diverse consequences that affect the outcome (e.g., the different energetic backgrounds of anoxia-induced and ouabain-induced SD will influence whether and how K_ATP_ channels are involved). Impairing barrier and ion homeostasis functions of the BBB predisposes towards SD generation without affecting the trajectory of individual SD events. The large investment of NKA and mitochondria in perineurial cells of the BBB (Hou et al., 2014; Hoyle, 1986; Smith and Shipley, 1990) indicates its importance for ion homeostasis of the CNS. It also suggests that this final common path of K^+^ ions out of the neuropil and into the hemolymph will be most vulnerable to cellular stress and the accumulation of damage. SD arises close to the BBB at a point of vulnerability and propagates laterally and more deeply into the neuropil. Once triggered, SD can mostly be attributed to transient, abrupt cell swelling (shrinkage of the restricted interstitium) that causes high [K^+^]_o_ and increased tissue resistivity. These processes interact with each other in a positive feedback manner (high [K^+^]_o_ causes cell swelling causes tissue resistivity increase causes high [K^+^]_o_). The nature of recovery from SD, or the failure to recover, is determined by the properties of continuous, background ion homeostatic mechanisms, such as NKA, or the accumulation of cellular damage.

In the presence of ouabain, the time to anoxic SD is immediately shortened from ∼8 mins to 1 − 2 mins and this latency does not change with repeated anoxic SDs during the course of TPP reversal (= NUP) over the following hour (Van Dusen et al., 2020). This intriguing observation might be explained if the initial effect of anoxia is to activate mitochondria through NKA (Baeza-Lehnert et al., 2019), generate more ATP and delay SD. Inhibition of NKA in perineurial cells of the BBB would prevent this. Given the dense packing of mitochondria in perineurial cells of ganglia (not connectives) (Hoyle, 1986; Smith and Shipley, 1990) and the high density of NKA in the perineurial sheath (Hou et al., 2014), we suggest that a fruitful research direction to explore would be a potential role of perineurial mitochondria and their interactions with NKA in triggering SD, perhaps by release of an endogenous SD activator (Andrew et al., 2022),. Another interesting ultrastructural feature of insect perineurial cells is the close association between the numerous mitochondria and membranes resembling smooth endoplasmic reticulum (Smith and Shipley, 1990). The authors speculate that the membranes may be infoldings of glial cell membrane to juxtapose NKA in the membranes with the mitochondria, consistent with the above suggestion. Another separate possibility is that the membranes are indeed endoplasmic reticulum. Mitochondrial endoplasmic reticulum contact (MERC) sites are important structures that facilitate Ca^++^ homeostasis and responses to metabolic changes (Giacomello and Pellegrini, 2016), and play an important role in the pathology of ischemic stroke (Zhang et al., 2025). Furthermore, RNAi knockdown of the *Drosophila* ortholog of Pdzd8, which is highly expressed in the fly CNS and is the ortholog of a mammalian MERC protein, increases lifespan and mitophagy suggesting that increased mitochondrial turnover supports the healthy aging of neurons (Hewitt et al., 2022). Finally, as noted above, pharmacological treatments are often limited by a lack of target-specificity and the difficulty of restricting the cell types exposed to the treatment. We strongly recommend that future research on insect models of SD, NUP and the BBB capitalize on the powerful genetic tools for time- and tissue-specific manipulation of identified molecular mechanisms of CNS ion homeostasis available for *Drosophila*.

## Acknowledgements

We thank Mads Andersen, David Andrew, Jens Dreier, and Heath MacMillan for their comments on a previous version of the manuscript. Funded by Discovery Grants # 04561-2017 to RMR and # 05841-2018 to AD from the Natural Sciences and Engineering Research Council of Canada.

## Abbreviations

BBB: blood brain barrier
CNS: central nervous system
DC: direct current
DIDS: 4,4ʹ-diisothiocyanostilbene-2,2ʹ-disulfonic acid
KCC: K^+^/Cl^−^ cotransporter
MDR: multidrug resistance
MERC: mitochondrial endoplasmic reticulum contact
MIA: monosodium iodoacetate
NACA: N-acetylcysteine amide
NKA: Na^+^/K^+^-ATPase
NKCC: Na^+^/K^+^/2Cl^−^cotransporter
NUP: negative ultraslow potential Pgp − phospho-glycoprotein
SD: spreading depolarization
SIET: scanning ion electrode technique
TTP: transperineurial potential
TPP30: transperineurial potential after 30 mins of ouabain exposure

## Notes

### Competing Interest Statement

The authors have declared no competing interest.

